# Plant phenotypes as trajectories: 38-yr monitoring reveals that shape of lifetime fecundity schedule is under selection in a long-lived shrub

**DOI:** 10.1101/2024.05.11.593670

**Authors:** Carlos M. Herrera

**Affiliations:** Estación Biológica de Doñana, Consejo Superior de Investigaciones Científicas (CSIC), Américo Vespucio 26, E-41092 Sevilla, Spain

**Keywords:** demography, individual variation, *Lavandula latifolia*, life history, lifetime fecundity schedule, plant phenotype, plant senescence, phenotypic selection

## Abstract

Defining a phenotype is sometimes problematic in the case of modularly-built, nonunitary organisms with indeterminate growth such as plants. This paper presents a proof of concept of the evolutionary significance of considering lifetime trajectories of individual plants as a component of their phenotypes. Size and inflorescence production were monitored for the whole reproductive lifespans of *N* = 128 individuals of the Mediterranean woody shrub *Lavandula latifolia* (Lamiaceae) over a 38-year period to address the following questions: Did individuals vary in lifetime trajectories of size and fecundity?, and Were parameters describing individual trajectories significant predictors of cumulative lifetime reproduction? Individuals differed widely in lifetime fecundity and in every parameter describing lifetime fecundity schedule (age at first and last reproduction, longevity, and mean, variance, skewness and kurtosis of lifetime temporal distribution of inflorescence production). Phenotypic selection analysis revealed significant relationships between parameters of individual lifetime schedules and a surrogate of relative fitness of individuals (cumulative lifetime production of inflorescences divided by the average for all individuals). Significant selection gradients involved positive and negative directional selection, as well as instances of nonlinear selection, which showed that plants with certain shapes of lifetime fecundity schedules had fitness advantage over others. The notion of plant phenotypes as trajectories was strongly supported by the combined findings that individuals differed greatly in their “appearances” with regard to the way in which size and fecundity unfolded over lifetime, and that selection efficaciously “saw” such variation.

## Introduction

Defining the phenotype of an organism is central to understanding the links between selection and potential evolutionary change, and how selection “sees” and “filters out” the variants of individual appearance that commonly coexist in natural populations of the same species. As stressed by Lewontin (1974, p. 19), “it is the evolution of the phenotype that interests us … [because] … what are ultimately to be explained are the myriad and subtle changes in size, shape, behavior, and interactions with other species that constitute the real stuff of evolution”. Selecting discrete or continuous measurable traits of individuals for obtaining a representation of their appearance is a trivial matter when one deals with unitary, non-modularly built organism with determinate growth such as most animals, but it is somewhat problematic in the case of modularly-built, nonunitary organisms with indeterminate growth such as most plants (Chitwood and Topp, 2015; Herrera, 2024). For example, defining the leaf or seed phenotype of an individual plant which can bear up to thousands of nonidentical copies of each of these kinds of homologous, reiterated structures poses a conceptual challenge, and it has been proposed that intraplant distribution in quantitative traits of reiterated structures should be considered a relevant dimension of its phenotype or appearance (Herrera, 2024).

One additional, insufficiently recognized difficulty to define plant phenotypes is that single-value traits of individuals (e.g., plant size), or features of the subindividually variable organs (e.g., leaf traits), can vary in the short term or over a plant’s lifetime due to ontogenetic processes (Herrera, 2009; Kulbaba et al., 2017; Steppe et al., 2011; Harder et al., 2019). In organisms with long life cycles such as perennial plants phenotypic traits will change with age, and development can be highly variable among individuals (Coleman et al., 1995; Chitwood and Topp, 2015). This points to the need of somehow incorporating a temporal dimension into the description of individual plant phenotypes for those traits that experience discernible ontogenetic changes over the plant’s lifetime (e.g., size, chemical, structural and functional traits of single leaves; Goodger et al., 2006; Greenwood et al., 2008; Steppe et al., 2011; Ji et al., 2021). Whenever individuals in a plant population differ in the properties of their lifetime trajectories for a given phenotypic trait, the *shape* of the trajectory will in itself be an element of the phenotype (my usage here of “phenotypes as trajectories” bears no relationship to the concept of “phenotypic trajectories” used to depict evolutionary changes of species or populations in phenotypic space; Adams and Collyer, 2009; Billman et al., 2014).

This paper presents a proof of concept of the evolutionary significance of considering lifetime trajectories as part of individual plants’ phenotypes. The study is based on data obtained by monitoring the growth and reproduction of a large sample of individuals of the Mediterranean woody shrub *Lavandula latifolia* (Lamiaceae) over their whole reproductive lifespans. The shape of lifetime reproductive schedule is taken here as a particular instance of phenotypic trajectory. The following specific questions will be addressed: (1) Did *L. latifolia* individuals differ substantially with regard to descriptive parameters of their lifetime size and fecundity trajectories, so that it makes sense to incorporate such trajectories into the description of their phenotypes ?; and (2) If individual differences in lifetime trajectory did exist, were descriptive parameters of individual trajectories significant predictors of cumulative lifetime reproduction, so that the existence of phenotypic selection on lifetime fecundity schedules can be inferredResults will provide affirmative answers to these two questions by showing that, when monitored over their entire reproductive lifetimes, individuals varied widely in size and fecundity trajectories, and that such variation was related to differential lifetime fecundity. Although some results presented here are also relevant in the light of plant demography and life history evolution (see Discussion), most of these aspects will not be considered in detail, as the main purpose of this article is to highlight the conceptual significance of considering the shape of lifetime trajectories as a phenotypic component in long-lived plants.

## Materials and methods

### Study plant

*Lavandula latifolia* is an evergreen shrub inhabiting forest clearings and edges in mid-elevation woodlands of the eastern Iberian Peninsula (see Fig. A1 in Appendix A of Herrera and Jovani, 2010, and Fig. 1 in Alonso et al., 2018, for photographs). The species reproduces exclusively by seeds, which are small (∼1 mg) and lack special mechanisms for dispersal. Plants have a main taproot and a short (usually <15 cm) main woody stem, and can be accurately aged by counting growth rings. Branching is dichasial, generally conforming to Leeuwenberg’s development model (a sympodial succession of equivalent sympodial units, each of which is orthotropic and determinate in its growth; Hallé et al., 1978; Hallé 1986). This branching pattern leads to crowns of adult plants being made up of distinct leaf clusters borne by short stems, many of which produce one terminal inflorescence in early summer. Each of these leaf clusters represents a “module” according to Hallé’s (1986, p. 78) definition (“the leafy axis in which the entire sequence of aerial differentiation is carried out, from the initiation of the meristem that builds up the axis to the sexual differentiation of its apex”). This produces an architecturally-mediated relationship between the number of inflorescences produced by an individual shrub at a given season and the total number of actively growing modules, as well as a close correlation between current leaf biomass and annual inflorescence production by individual plants. In the population studied, yearly seed production by adult plants was predicted by the number of inflorescences (*R*^2^ = 0.44, Herrera, 1991), and individual variation in average per-seed viability (emergence rate and seedling survival) was comparatively narrow in relation to the broad differences in inflorescence production (Herrera, 2000). Annual inflorescence production can therefore be used as a reasonable proxy for fecundity comparisons both across individuals and across years within individuals. Further details on natural history, reproductive biology and demography of *L. latifolia* in the study area can be found in Herrera (1991), Herrera and Jovani (2010), Herrera and Bazaga (2016), Alonso et al. (2018) and Herrera et al. (2021).

**Figure 1.**
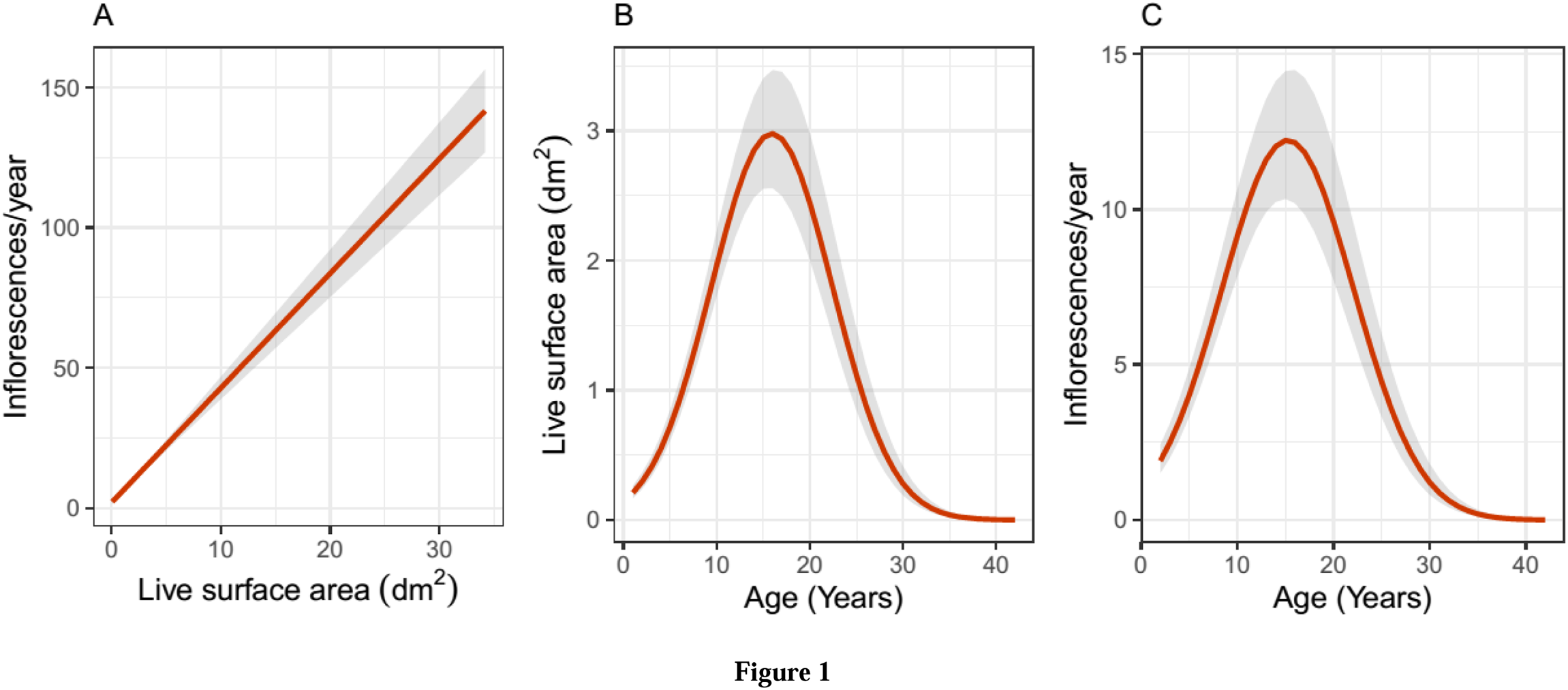
Model-predicted, average population-level estimates of the relationships linking current plant size with yearly inflorescence production (A), and plant age with size (B) and yearly inflorescence production (C), over the lifetime of individual plants. Predictions were obtained by fitting random-slopes, random-intercepts linear mixed-effect models to the data in which the variable on the vertical axis was the response variable, the variable on the horizontal axis was the fixed-effect predictor (the quadratic transform was also included in the case of B and C), and plant was included as random factor. In B and C the response variable was log-transformed for the analyses, and model predictions were back-transformed to the original scale of measurement. Grey areas represent standard errors around estimates.

### Field methods

This study was conducted during 1986-2023 at the “Aguaderillos-2” site of Herrera (1988, 1991), located at 1220 m elevation in a mixed woodland of *Pinus nigra* and *Quercus rotundifolia* in the Sierra de Cazorla, Jaén province, southeastern Spain (site coordinates 37.9612°N, 2.8847°W). A permanent 200-m^2^ plot was established there in July 1986 which remained undisturbed for the whole duration of this study. In August 1986 all *L. latifolia* plants within the plot bearing ≥10 leaves (roughly ≥ 2 yr in age) were mapped, tagged and measured (see below), and the number of current year’s inflorescences counted. Only the subset of plants which at the time of tagging in 1986 were nonreproductive juveniles and reproduced in two or more years over 1987-2023 will be considered here (*N* = 128). All these plants had already died by the summer of 2023, hence the data used here encompass their complete reproductive lifespans. All plants were checked every summer (August-September) between 1987-2023. On each occasion, individuals found dead were digged up, collected, and their age at the time of death determined by ring counting. For each individual that remained alive, the number of inflorescences produced in the current year was counted and, on the even years, their size was assessed by measuring the two major axes of the canopy and then computing the area of the horizontal projection by assimilating the shape to an ellipse. In senescent plants with discontinuous canopies comprising two or more living sectors (see photographs in Fig. A4, Appendix A of Herrera and Jovani, 2010, for examples), measurements were taken and areas computed separately for each living sector, and then summed up to obtain a single size estimate for the individual.

### Data analysis

The birth year of every monitored individual was determined by substracting the age at death from the year of death, and this information was then used to retrospectively compute the age of every plant in each study year. These data were then merged with those on yearly fecundity (number of inflorescences) and size measurements (estimated canopy area) to obtain individual lifetime profiles of size and fecundity in relation to current age. As will be shown in the results, size and fecundity lifetime profiles were closely correlated, and only those for fecundity will be analyzed here in detail.

The lifetime fecundity schedule of each plant was characterized quantitatively by combining two sets of parameters, hereafter named “landmark” and “shape” lifetime parameters. The landmark parameters refer to individual life history attributes from the classical “generalized triangular reproductive function” (Lewontin, 1965; Harper, 1977): age at first reproduction, age at last reproduction, and age at death (“total lifespan” hereafter). The shape parameters summarize numerically the individual trajectories by taking advantage of the fact that, when complete lifetime data are used, individual curves of yearly fecundity versus age actually represent frequency distributions of the age at which reproductive units (inflorescences in the present instance) were produced. The first four statistical moments of such frequency distributions (mean, variance, skewness, kurtosis) were computed for each plant and provided the shape parameters of lifetime fecundity schedules. The intensity of linear (directional, *β*_i_) and quadratic (stabilizing or disruptive, *γ*_i_) selection gradients on lifetime fecundity schedules was investigated by adopting a phenotypic selection approach, based on regressing total inflorescences produced over individual lifetime against landmark and shape parameters and their quadratic transforms (Lande and Arnold, 1983; Svensson, 2023).

All statistical analyses reported in this paper were carried out using the R environment (R Core Team, 2023). Average population-level trajectories of plant size and annual inflorescence production in relation to plant age were estimated by fitting random-slopes, random-intercepts linear mixed-effect models to plant area and inflorescence number data (both log-transformed), with age and age-squared as fixed-effect predictors and individual plant as random factor. The population average for the within-plant relationship over lifetime between plant size and inflorescence production was evaluated by fitting a random-slopes, random-intercepts linear mixed-effect model to the data with plant size as fixed-effect predictor and plant as random factor. The function lmer in the package LME4 (Bates et al., 2015) was used for fitting mixed-effect models, and predicted shapes of the relationships and associated confidence intervals were obtained by applying to each fitted model the function ggpredict from the GGEFFECTS package (Lüdecke, 2018). Marginal *R*^2^ (variance explained by fixed factors) and conditional *R*^2^ (variance explained by both fixed and random factors) of mixed-effect models (Nakagawa and Schielzeth, 2013) were obtained with function r2_nakagawa in the PERFORMANCE package (Lüdecke et al., 2021).

Phenotypic selection on descriptive parameters of individual lifetime fecundity schedule was assessed by fitting a linear model to the data using function lm in the STATS package. For the estimation of selection coefficients (*β*_i_, *γ*_i_) and their standard errors, the response variable was the cumulative production of inflorescences over each individual’s lifetime divided by the average of this value for all individuals in the sample (“relative fitness”, Lande and Arnold, 1983). The predictors were the three landmark and four shape schedule parameters defined above plus their quadratic transforms, all of which were included simultaneously in a single multiple regression model. All variables in the model except the response were scaled to mean zero and standard deviation unity, so that the estimated model parameters represented standardized estimates (Lande and Arnold, 1983). Individual lifetime fecundity departed substantially from normality, which invalidated asymptotical statistical significance tests of parameters based on their standard errors from this model. To evaluate the statistical significance of predictors, the model was re-run after transforming fecundity values logarithmically, which satisfactorily normalized residuals, and significance of parameter estimates (i.e., *β*_i_ or *γ*_i_ ≠ 0) was then evaluated with ordinary *F* tests using the Anova function in the car package (Fox and Weisberg, 2019).

## Results

### Population-level relationships between size, fecundity and age

After statistically accounting for individual heterogeneity by including plant as a random factor in the model, there existed a close linear relationship at the population level between yearly inflorescence production and current plant size (area alive) over an individual plant’s lifetime (Figure 1A; Chi-square = 316.14, df = 1, *P* < 1e-06). The model fit well the data, and the rough similarity between marginal (0.598) and conditional (0.788) *R*^2^ values suggests that individual variation in the size-fecundity relationship was only a secondary source of variance in yearly inflorescence production.

As would be anticipated from the close relationship between size and fecundity over individual plant lifetime, the model-estimated, population-level relationship linking plant size (Figure 1B; marginal *R*^2^ = 0.353, conditional *R*^2^ = 0.852) and yearly inflorescence production (Figure 1C; marginal *R*^2^ = 0.237, conditional *R*^2^ = 0.820) with plant age were virtually identical. On average, both size and yearly inflorescence production first increased steadily until plants reached ∼15 years and then declined steadily until death. The substantial difference between marginal and conditional *R*^2^ in each of these two models provides an indirect indication that the among-plant differences subsumed into the random terms were substantial and contributed most variance to the response variables in the models, as detailed in the following section. Given the close similarity between lifetime patterns for size and inflorescence production, only the latter will be considered hereafter.

### Individual lifetime fecundity schedules and parameters

The lifetime reproductive schedules of all *L. latifolia* plants studied are plotted side by side in Figure 2 to graphically highlight the extraordinary multivariate diversity of individual lifetime fecundity trajectories. Individual schedules included, for example, broad-flat ones with slowly increasing production of inflorescences over all the reproductive lifespan; broad-weakly convex ones denoting long reproductive lifespans with a shallow peak at some intermediate age; or narrow-convex shapes with reproduction concentrated over short reproductive spans. Schedule diversity involved variation in age at first reproduction (range = 2-17 years, interquartile range IQR = 5-7), age at last reproduction (range = 6-42 years, IQR = 18-24), and total lifespan (range 10-44 years, IQR = 20-26). Across plants, age at first reproduction was not significantly correlated with either age at last reproduction or total lifespan (*r*_s_ = 0.106 and 0.144, *P* = 0.23 and 0.11, respectively, *N* = 128, Spearman rank correlation). The extensive individual variation in shape of lifetime fecundity schedules (Figure 2) was also reflected in the broad frequency distributions of the four shape parameters considered (Figure 3).

**Figure 2.**
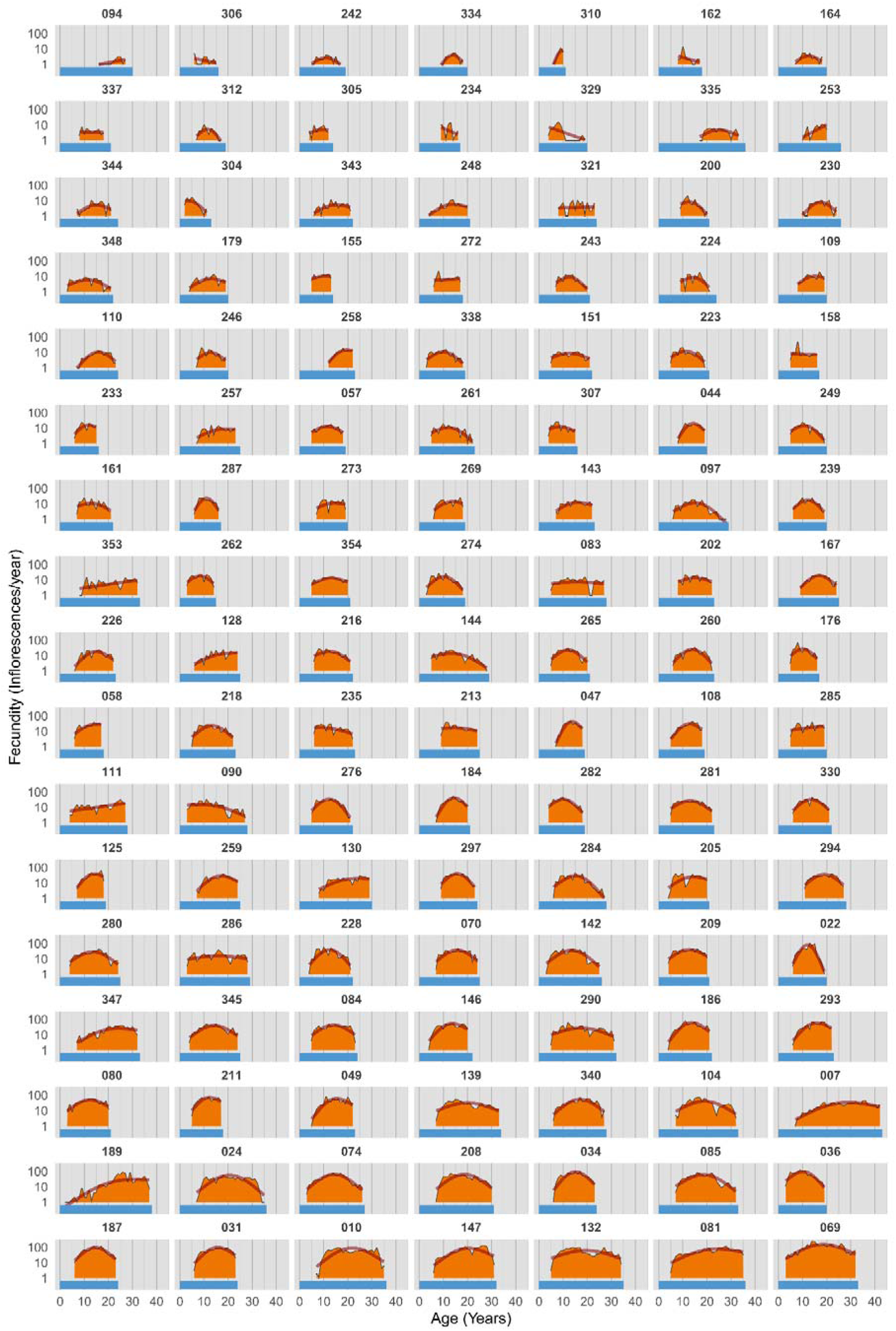
Lifetime reproductive schedules of individual *Lavandula latifolia* plants that were monitored for their whole reproductive lifespans (only the *N* = 119 plants in the sample that flowered on at least 5 years are shown). Each panel corresponds to a different individual (identified by its field tag), and shows lifetime variation in yearly inflorescence production (orange-filled area under thin black lines), smoothed lifetime fecundity trajectory (thick orange line, fitted to yearly data using a gam smoother), and total lifespan (horizontal blue bar). Plants are ordered by increasing lifetime fecundity (orange area, from left to right, and top to bottom). Note logarithmic scale on vertical axis.

**Figure 3.**
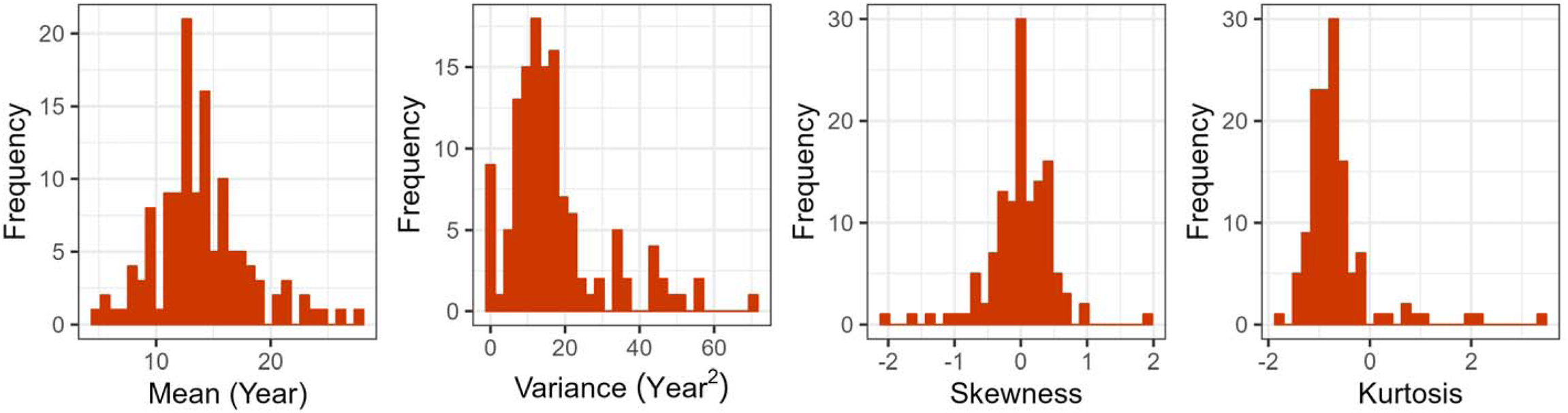
Frequency distributions of the four shape parameters describing individual lifetime fecundity curves in the set of *Lavandula latifolia* plants studied (*N* = 128). Shape parameters are the four moments of the frequency distributions of the ages at which lifetime inflorescences were produced by each plant (mean, variance, skewnes and kurtosis).

The diversity of lifetime fecundity schedules represented in the set of monitored plants is well synthesized by the broad two-dimensional scatter of the empirical cumulative distributions of fecundity against age when data points for all individuals are plotted together (Figure 4). The cumulative proportion of inflorescences produced up to a given age, or the age required to achieve a given proportion of cumulative lifetime fecundity, were both extremely variable among plants. For example, some individuals had already produced 50% of their eventual lifetime inflorescences before being 10 years old, while others only achieved that cumulative fecundity threshold at ages ≥20. At age 20 some plants had produced <25% of lifetime inflorescences while many had already produced >75%.

**Figure 4.**
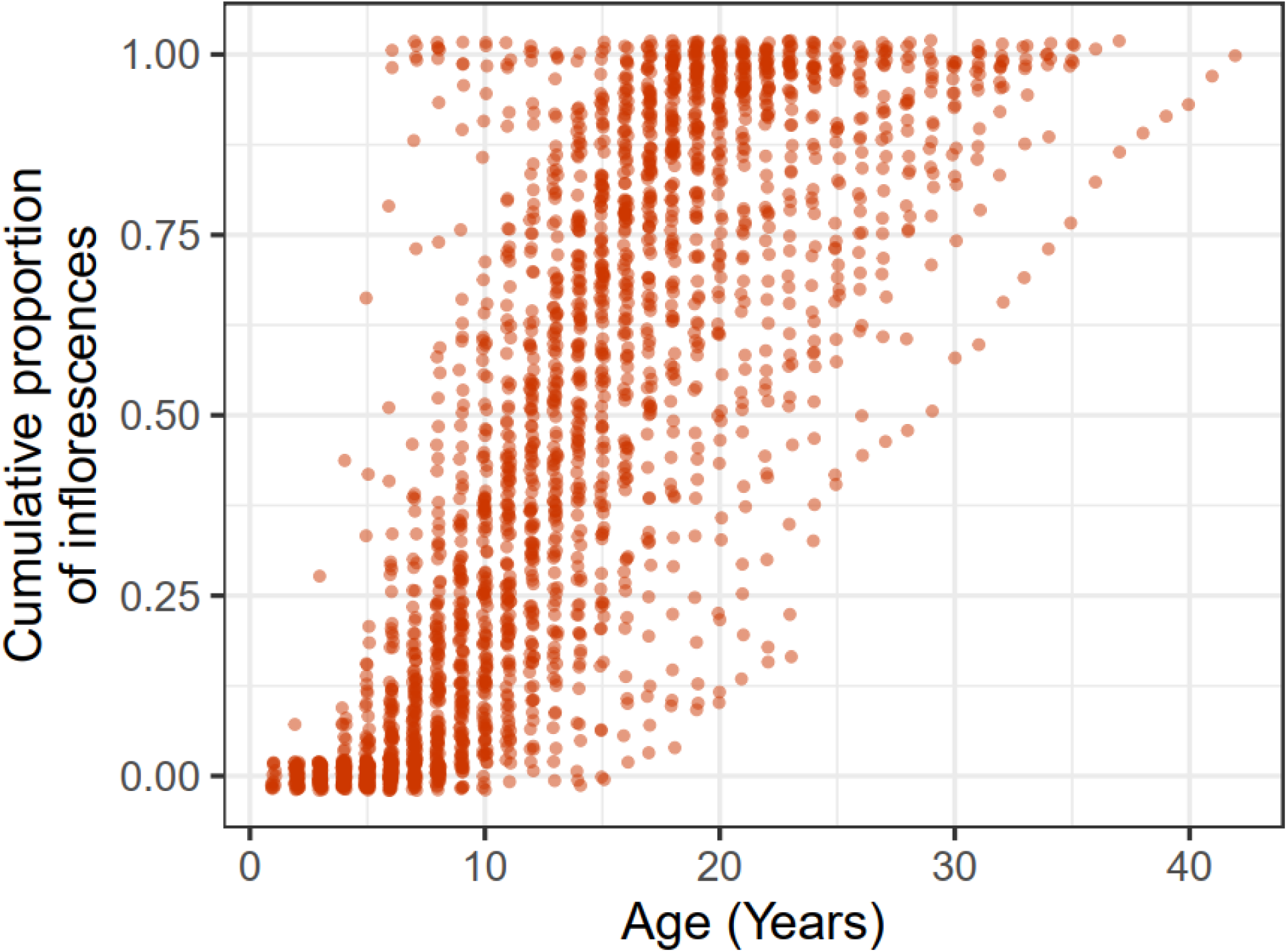
Cumulative density distributions of the number of inflorescences produced over lifetime by individual *Lavandula latifolia* plants. To avoid cluttering and highlight the broad individual variability represented in the sample of plants studied, lines connecting points for individual plants are omitted. For any given age, each point corresponds to a different plant, and the vertical scatter illustrates individual variation in the proportion of total lifetime inflorescences that were produced up to and including that age. A small random noise has been added to the horizontal and vertical coordinates of data points.

### Phenotypic selection on lifetime fecundity schedule

Individual plants differed widely in lifetime fecundity (range = 3-2810 inflorescences, IQR = 86-406). After transformation to individual relative fitness, lifetime fecundity was significantly predicted by the landmark and shape parameters of lifetime fecundity schedules (*F*_14, 113_ = 30.6, *P* < 2.2e-06, adjusted *R*^2^ = 0.791; multiple regression of log-transformed relative fitness against standardized landmark and shape parameters). Each descriptive parameter of the fecundity schedule was a significant predictor of lifetime inflorescence production after statistically controlling for the combined effects of the rest of parameters (Table 1). In the case of landmark parameters only the linear terms had significant effects on lifetime fecundity: age at last reproduction was directly related to lifetime fecundity, while age at first reproduction and total lifespan were both inversely related. These relationships denote that plants starting to reproduce earlier in life, discontinuing reproduction at a later age, and/or having shorter longevities tended to be at relative fecundity advantage. In the case of shape parameters there were both linear and quadratic significant effects. Lifetime schedule curves with lower variance and kurtosis (i.e., narrower curves) were at relative fecundity advantage. Significant quadratic effects had either positive (variance, skewness) or negative (mean) signs (Table 1), revealing instances of stabilizing and disruptive selection, respectively.

**Table 1.** Standardized linear and quadratic selection gradients on the seven parameters used to describe the lifetime fecundity schedules of *Lavandula latifolia* plants studied (*N* = 128). Statistical significant terms (*P* < 0.05) are shown in bold type.

| Fecundity schedule parameter | Terms in linear model |  |  |  |  |  |
| --- | --- | --- | --- | --- | --- | --- |
|  | Linear |  |  | Quadratic |  |  |
| | $\beta$ | SE | $P$-value | $\gamma$ | SE | $P$-value |
| Landmark parameters: |  |  |  |  |  |  |
| Age at first reproduction | **-1.439** | 0.237 | **2.8E-08** | +0.192 | 0.049 | 0.32 |
| Age at last reproduction | **+1.442** | 0.994 | **4.2E-05** | +0.848 | 0.364 | 0.50 |
| Total lifespan | **-0.512** | 0.778 | **0.012** | -0.253 | 0.370 | 0.53 |
| Shape parameters: |  |  |  |  |  |  |
| Mean | +1.525 | 0.485 | 0.11 | **-0.665** | 0.126 | **0.024** |
| Variance | **-1.661** | 0.469 | **5.5E-05** | **+0.251** | 0.098 | **0.0051** |
| Skewness | +0.216 | 0.149 | 0.78 | **+0.287** | 0.115 | **0.022** |
| Kurtosis | **-0.582** | 0.226 | **0.0059** | -0.029 | 0.048 | 0.18 |

## Discussion

### Population average versus individual lifetime schedules

Monitoring the whole reproductive lifespans of a large sample of *L. latifolia* plants of known birth date has revealed a well-defined average lifetime fecundity schedule at the population level. It involved a steep increase in size and reproduction up to reaching a peak about midway in lifetime (∼15 years), followed by a steep decline ending up in the plant’s death (∼30-40 years). This two-stage lifetime pattern was parsimoniously accounted for by a shift in the dynamic balance between module formation and module death, in accordance with the notion of the plant individual as a population of semi-autonomous modules (White, 1979; Watkinson and White, 1985). During the first phase the number of living modules and inflorescences produced annually increased steadily with age, while a senescence period (meant here as age-dependent decline in size and reproduction) ensued during the second phase. In this period, inflorescence production declined and living area shrank because yearly production of new modules did not compensate for the increasing annual mortality of modules. The present results for *L. latifolia* (see also Herrera and Jovani, 2010) contribute to the accumulating evidence suggesting that gradual individual senescence is probably more frequent in natural populations of perennial plants than previously thought (Silvertown et al., 2001; Munné-Bosch, 2008, 2015; Roach and Smith, 2020).

The population average for the lifetime fecundity schedule was only weakly representative of the broad range of individual schedules actually occurring in the sample of *L. latifolia* plants studied, whose general shapes, temporal landmarks and shape descriptors varied widely among individuals. The lifetime fecundity schedule represents a keystone component in the demography of populations and the evolution of life history traits (Lewontin, 1965; Harper, 1977; Stearns, 1992), yet I was unable to locate any published research reporting individual lifetime fecundity schedules for woody plants. Instances of broad individual variability in supra-annual patterns of fruit production have been frequently reported for tropical and temperate woody plants (e.g., Janzen, 1978; Crawley and Long, 1995; Herrera, 1998; Munilla and Guitián, 2014, Pérez-Ramos et al., 2014), but these data are poor indicators of individual lifetime schedules because they ordinarily refer to plants of unknown age and comprise periods much shorter than the presumed reproductive lifespan. It is not possible therefore to know the extent to which the large individual variability found here for *L. latifolia* is a general characteristic of individual lifetime schedules of woody plants.

The practical difficulties inherent to collecting the data needed for elucidating individual fecundity schedules of long-lived plants have been emphasized many times (Harper and White, 1974; Janzen, 1978; Clarke, 1992; Salguero-Gómez et al., 2015). For example, the modal duration of studies included in the initial version of the comprehensive COMPADRE database on plant demography was “4 years, corresponding to the length of an average PhD project, as well as that of most funding agencies” (Salguero-Gómez et al. 2015, p. 213). The modal duration of studies subsequently declined to 3 years in the most recent version (COMPADRE 6.23.5.0, accessed 8 March 2024), where only 0.85% of the studies included (*N* = 8823) were on trees/shrubs and had a duration ≥25 years. Since 62% of the angiosperms are woody species (Zanne et al., 2014), their underrepresentation in demographic studies speaks of the practical difficulties noted above and denotes a considerable bias in the demographic information available for plants. Statistical methods devised to circumvent limitations imposed by unknown plant age and fragmentary fecundity data can successfully estimate average lifetime schedules and other demographic parameters at the population or species levels (Clarke, 1992; Silvertown et al., 2001; Jones et al., 2014; Wenk and Falster, 2015; Caswell, 2019). These estimates are valid for comparative purposes (Silvertown et al., 2001; Jones et al., 2014; Wenk and Falster, 2015; Qiu et al., 2021), but the lack of empirical information on the magnitude of individual variation in lifetime parameters prevents formal statistical testing of differences between species or populations, since uncertainty around average estimates cannot be evaluated (van Daalen and Caswell, 2017). Most importantly, average schedules for groups of individuals do not allow for tests of the relationships between individual fitness and features of lifetime fecundity schedules which have been historically hypothesized to underly life history evolution (see references in Introduction).

### Selection on lifetime fecundity schedule

The phenotypic selection analysis revealed strong, significant relationships between all landmark and shape parameters of individual lifetime schedules and the surrogate used here to estimate the relative fitness of *L. latifolia* individuals (cumulative lifetime production of inflorescences divided by the average for all individuals). Selection gradients on landmark parameters involved positive directional selection on age at last reproduction, and negative directional selection on age at first reproduction and total lifespan, which revealed that individuals combining early start of reproduction, long reproductive lifespan, and short total lifespan were at relative fecundity advantage. Selection on shape parameters involved both directional (negative on variance and kurtosis) and quadratic (stabilizing on the mean, disruptive on variance and kurtosis) components, again reflecting that everything else being equal certain shapes of lifetime fecundity schedules had relative fitness advantage over others. Selection on the set of descriptive parameters of lifetime schedules was consistently strong, as revealed by the high absolute values of *β*_i_ (median |*β*_i_| = 1.44, range = 0.22-1.66) and *γ*_i_ (median |*γ*_i_| = 0.25, range = 0.03-0.85). These figures fall in the uppermost quantiles of the distributions of (absolute) selection gradients previously reported for all types of traits in natural or seminatural plant populations, including life history features (see Kingsolver et al., 2001; Siepielski et al., 2009; Kingsolver and Diamond, 2011; Caruso et al. 2018, 2020, for comprehensive reviews of phenotypic selection studies). It is not possible to elucidate whether the higher selection gradients found here in relation to most previous phenotypic selection studies stem from the type of traits considered, the lifetime nature of the fitness surrogate data (lack of estimates of lifetime fitness is “a major fault of most studies of natural selection“; Endler, 1986, p. 162), or some combination of both.

The usual caveats raised on phenotypic selection approaches, particularly those questioning inferences of causality drawn from trait-fitness relationships (Svensson, 2023), apply also to the present study, since it is not possible to identify the proximate selective mechanism(s) accounting for the strong selection observed on landmark and shape parameters of individual lifetime fecundity schedules. Furthermore, the phenotypic selection analysis conducted here was based on a proxy of fecundity rather than on actual fecundity data (i.e., number of offspring produced). Keeping these caveats in mind, it is worth noting that results of this study are among the few instances to date of empirical support to some central tenets of life history evolution theory in a *within-population* context. Regarding landmark parameters, the strong negative directional selection on age at first reproduction found here agrees with the selective value consistently attached to reproductive precocity in models of life history evolution, particularly in the case of species with colonizing, expanding populations (Cole, 1954; Lewontin, 1965; Stearns, 1976, 1992; Harper, 1977) such as *L. latifolia* and, more generally, Mediterranean woody plants whose populations are exposed to recurrent wildfires (Guiote and Pausas, 2023). Positive directional selection on reproductive lifespan also agrees with some assumptions of life history theory (Stearns, 1992). Shape parameters of lifetime fecundity schedules have been rarely considered in theoretical formulations of life history evolution, but selection on mean, variance and skewness found here are implicit in some of the hypothesized scenarios envisaged by Lewontin (1965) under his “generalized triangular reproductive function”.

### Phenotypes as trajectories

The significance of ontogenetic changes in long-lived plants has been previously considered in a variety of ecological and evolutionary contexts, but there have been few attempts at placing the phenomenon in the conceptual framework of an expanded definition of plant phenotypes (but see, e.g., Harder et al., 2004; Herrera, 2009; Chitwood and Topp, 2015; Kulbaba et al., 2017; Harder et al., 2019; Herrera et al., 2021; Barton, 2024). The set of landmark and shape features used here to characterize the lifetime schedules of growth and reproduction of individual *L. latifolia* plants represent a description of the way in which individual genotypes unfolded over its lifetime with regard to two important facets of its external (i.e., phenotypic) appearance, namely plant size and annual number of inflorescences. The heterogeneous assortment of shapes depicted in Figure 2 portrays an assembly of “visible and measurable appearances” (Wanscher, 1975, p. 144) of monitored plants, and hence are genuine elements of their phenotypes. Further support to the notion of phenotypes as trajectories was provided by the finding that selection was somehow able to “see” the variation existing among individuals, so that these individual differences were eventually related to lifetime fecundity. The possible evolutionary significance of selection on phenotypic trajectories will obviously depend on whether the causal mechanisms responsible for individual differences in lifetime schedules have some heritable basis (van Daalen and Caswell, 2017). Given the considerable evidence supporting a role of genetic and (heritable) epigenetic control on individual development, growth and reproduction in plants (Bogan et al, 2024; Mitchell_Olds, 1996a,b; Turck and Coupland, 2013; Kottler et al., 2018; Herrera et al., 2019), and the considerable genetic and epigenetic diversity exhibited by the marked population of *L. latifolia* studied (Herrera and Bazaga, 2016), it seems plausible to speculate that observed individual differences in lifetime trajectories were at least in part due to heritable factors. Unfortunately, the long time required for estimating the heritability of lifetime schedule parameters using quantitative genetics methods based on controlled crosses and parent-offspring correlations will in fact render such heritability estimates unknowable for most woody plants.

While the concepts of gene and genotype have been a recurrent topic of discussion throughout the history of genetics (Roll-Hansen, 2014), the description and use of the closely associated concept of phenotype have remained remarkably stable and undebated since its inception more than a century ago (Wanscher, 1975; Herrera, 2024). There are reasons, however, for revising the concept of phenotype in the case of modularly-built, nonunitary organisms with indeterminate growth such as plants and some colonial animals (Chitwood and Topp, 2015; Herrera, 2024). On one side, there is the need of accommodating instantaneous intraindividual variability in features of reiterated organs into the concept of phenotype by treating such variation as trait distributions rather than a simple average (Herrera, 2024). On the other hand, plants that grow, develop and reproduce over long periods demand the incorporation of individual lifetime trajectories as a critical temporal dimension of their respective phenotypes, as motivated in the Introduction to this article and supported by the results presented. Understanding the genotype-phenotype relationship was a major goal of the research agenda in genetics and evolutionary biology during the last century (Sturtevant, 1965; Lewontin, 1974; Falconer, 1981; Noble, 2015). This research tradition paid most attention to unraveling the genetic underpinnings of phenotypic traits and the mechanisms whereby genetic background accounted for the organisms’ appearance. In contrast, little effort has been historically devoted to scrutinize whether the original Johannsenian definition of phenotype (Johannsen 1909, 1911) suited equally well to all kinds of organisms or, alternatively, should in certain groups be superseded by a more elaborate concept (Herrera, 2024, and present study). An updated definition of phenotype should incorporate the ontogenetic, developmental and architectural peculiarities of organisms, particularly the multiplicity of nonidentical structures produced by single genotypes and the protracted period of steady phenotypic unfolding arising from ontogenetic shifts, growth schedules and internal modular dynamics.

## Acknowledgments

People helping with plot maintenance and plant monitoring over nearly four decades, or providing ideas and discussion on this project, are too numerous to remember or mention individually. Among them, special thanks are due to Manolo Carrión (†), Pedro Jordano, Roger Jovani, Luis López-Soria and Dori Ramírez. I am indebted to Conchita Alonso, Mónica Medrano and Juli Pausas for valuable suggestions which significantly improved an earlier version of the manuscript. Consejería de Medio Ambiente, Junta de Andalucía, provided permission to work in Sierra de Cazorla along with invaluable facilities. While writing this article I benefitted from grants from Ministerio de Ciencia, Innovación y Universidades (PID2022-141530NB-C22, DISTEPIC) and Generalitat de Valencia (PROMETEO-2021/040, FocScales).

## Data and Code Accessibility Statement

All the raw data and scripts required to replicate the analyses reported in this study will be deposited in figshare upon manuscript acceptance.

